# Structure and Variation in Dam–Daughter Bonds in Captive Rhesus Macaques

**DOI:** 10.64898/2026.05.27.728159

**Authors:** Brianne Beisner, Kelly F. Ethun

**Affiliations:** Division of Developmental and Cognitive Neuroscience, Emory National Primate Research Center, Emory University, Atlanta, GA 30329; Division of Animal Resources, Emory National Primate Research Center, Emory University, Atlanta, GA 30329; Department of Pathology and Laboratory Medicine, Emory School of Medicine, Emory University, Atlanta, GA 30322

**Keywords:** Nonhuman primates, cercopithecines, social instability, group management

## Abstract

Mother–daughter relationships are central to social organization in rhesus macaques, providing the foundation for kin familiarity, affiliative bias, and intergenerational rank inheritance. Understanding how these bonds are structured is therefore important for identifying the processes that contribute to matrilineal cohesion and for anticipating risks of social instability and matriline fragmentation in captive groups.

We studied 141 dam–daughter dyads in large, captive groups of rhesus macaques, using generalized linear models to examine associations between demographic variables and affiliative behaviors, and a multivariate clustering approach to characterize how these behaviors combine into distinct relationship types.

Our analyses identified eight distinct dam–daughter relationship types defined by combinations of three behavioral domains: grooming, spatial closeness, and agonistic support. Generalized linear models showed that demographic variables had domain-specific effects, with family size, age gaps, and dam age differentially associated with grooming, proximity, and support. These patterns were reflected in the clustering analysis, in which dyads differed not only in overall levels of behavior but also in how behaviors combined within dyads. For example, some dyads exhibited high grooming and support despite low spatial proximity, whereas others showed consistently high expression across all domains.

Together, these results demonstrate that dam–daughter bonds are multidimensional and structured through the combination of multiple behavioral domains influenced by demographic and life-history factors. Because these dyads scaffold interactions among close kin, variation in dam–daughter bond structure may influence the development and stability of broader matrilineal relationships. From an applied perspective, considering multiple behavioral domains when assessing social relationships can help identify dyads that may warrant closer monitoring and inform management strategies to support matrilineal cohesion and reduce the risk of social instability in captive groups.

## 1 Introduction

Housing social animals in captivity requires careful consideration of species-typical social structure to promote positive welfare. For instance, captive populations of rhesus macaques bred for biomedical research are often housed in large outdoor groups because they mimic key features of free-ranging groups (Hannibal et al., 2016). Such environments are associated with improved reproduction, psychological well-being, and translational (Haertel et al., 2023; Hannibal et al., 2016). Multigenerational structure in particular has been found to improve female reproductive success and offspring survival in captive rhesus macaque populations (Rox et al., 2022).

Female rhesus macaques live in matrilineally structured groups in which kin-based alliances support rank inheritance and matrilineally structured dominance hierarchies (Bernstein and Gordon, 1974; Sade, 1972, 1967). Captive multigenerational housing aims to preserve these dynamics by allowing females to remain in their natal groups. Because bonds among related females stabilize daily social interactions and dominance relationships (McCowan et al., 2017), understanding how these bonds form and are maintained has direct implications for the management and welfare of captive populations.

Matriline fragmentation poses a significant welfare concern in captive rhesus groups (Beisner et al., 2011; Ehardt and Bernstein, 1986; McCowan et al., 2017; Oates-O’Brien et al., 2010). Disruption of kin-based networks can destabilize dominance relationships, increasing the risk of severe aggression, social overthrows, and associated injury. Fragmented matrilines exhibit reduced grooming network cohesion and elevated conflict among related females (Beisner et al., 2026, 2011). However, prior work assessed cohesion at the family level, using a single behavioral domain (grooming) with less attention to dyadic processes (Beisner et al., 2011). Thus, while fragmentation can be detected, the specific relationships that generate or maintain cohesion (i.e., potential targets for management) remain unclear. Yet, kin bias in cercopithecine primates is expressed across multiple behavioral domains, including grooming, spatial proximity, and agonistic support (Kapsalis, 2004), each of which contributes to the formation and maintenance of social relationships and dominance structure. If these behaviors do not vary in concert, then cohesion cannot be fully captured by any single measure.

Here, we define matrilineal cohesion as the coordinated patterning of multiple affiliative behaviors among related females and examine it at its presumed foundational unit: the mother–adult daughter relationship. Mother–daughter relationships are among the strongest, most consequential kinship bonds. They scaffold familiarity and proximity among close kin, supporting kin recognition, affiliative bias, and the transmission of social bonds across generations (Berman and Kapsalis, 1999; Berman, 2004; Rendall, 2004). Sisters become familiar through shared association with their mother, and mothers provide agonistic support that facilitates intergenerational rank inheritance (Chapais, 1992, 1988). Thus, family cohesion is not automatic but constructed through multiple interaction domains.

Since mother-daughter bonds are foundational, their structure warrants careful consideration. Hinde (1976) emphasized that relationships are defined by behavioral patterning across domains over time rather than by single behaviors. Accordingly, mother–daughter relationships should be examined as multidimensional configurations (e.g., grooming, proximity, agonistic support) that vary across life-history stages. Furthermore, matriline cohesion cannot be fully inferred from single behaviors but instead emerges from integrated interaction patterns among kin.

Affiliative bonds in nonhuman primates have been quantified in a variety of ways. Many studies use rates of a single behavior, like grooming, to measure relationship strength. Others combine multiple behaviors into composite measures such as the dyadic or composite sociality index (Silk et al., 2010, 2009, 2003), or construct social networks based on a single affiliative behavior (Balasubramaniam et al., 2018, 2016; Vandeleest et al., 2025). Work on kin bias similarly draws on grooming, proximity, and agonistic support as indicators of preferential social investment (Berman, 2004; Kapsalis, 2004). Across approaches, behaviors are often analyzed independently or aggregated, with limited attention to how different domains integrate within dyads. However, recent findings demonstrate that relationships characterized by multiple coordinated affiliative behaviors are associated with improved physiological outcomes, whereas those expressed through a single behavioral domain (i.e., grooming) are not (Vandeleest et al., 2025). These findings suggest that similar rates of one behavior can mask variation in multidimensional relationship structure. A central question, therefore, is how multiple behavioral domains combine within mother–daughter dyads to generate the relational architecture of matrilines.

This study examines how demographic factors shape variation in mother–daughter relationships in rhesus macaques, asking whether these variables uniformly strengthen or weaken affiliative and agonistic behaviors or instead constrain dyads to express affiliation through a limited set of multivariate relationship types. To address this, we combine behavior-specific generalized linear models with an unsupervised clustering approach that captures the joint structure of multiple behavioral domains.

## 2 Methods

### 2.1 Study Site and Subjects

Subjects were rhesus macaques (*Macaca mulatta*) from five breeding groups housed at the ENPRC Field Station (Lawrenceville, GA). Group size ranged from 62 – 168 individuals. All groups were housed in indoor–outdoor compounds (outdoor: 232–1,450 m²; indoor: 15–35 m²). Animals had continuous access to drinking water and were fed a commercial diet *ad libitum* via automated feeders (Johnston et al., 2020). Diet was supplemented with produce and environmental enrichment (climbing structures, foraging devices, and manipulanda). All animals were SPF (free of SIV, simian T-lymphotropic virus, simian type D retroviruses, and herpes simian B virus). Behavioral data were collected between July 2022 and June 2025.

A total of 141 unique dam–adult daughter dyads were identified across the five study groups (17–41 dyads per group). Both dams and daughters were required to be present for at least 50% of observation sessions to be included in analyses.

### 2.2 Behavioral Data Collection

Behavioral observations were conducted by three trained observers who recorded affiliative and agonistic interactions among individuals aged ≥3 years. Groups were observed 12 hours per week for 10 months, yielding 480–528 observation hours per group. Affiliative behaviors were recorded using scan sampling and included grooming, huddling (all forms of body contact between two individuals), and proximity (within arm’s reach). Agonistic interactions were recorded using an all-occurrences event sampling protocol, including participant identity and associated aggressive (i.e., threats, lunges, chases, bites, and grappling or pinning) and submissive (i.e., freeze/turn away, move away, run away, silent bared teeth display, and scream) behaviors. Third-party agonistic interventions were also recorded. Support directed toward the victim of aggression was classified as aid-for-victim (AFV), whereas siding with the aggressor was classified as new-join-with (NJW). Data were recorded using the HandBase application on handheld devices or by voice recorder and later transcribed. As measured by Krippendorff’s alpha, inter-observer reliability was confirmed to be ≥97% accurate for individual identification and ≥88% reliability for behavioral coding.

### 2.3 Behavioral Outcome Variables

To measure the nature and the relative strength of dam-daughter relationships, we examined both affiliative interactions (huddle, proximity, and grooming) and agonistic support interactions (aid-for-victim). These behaviors are well established indicators of strong social bonds, particularly among kin. For each dam–adult daughter dyad, behavioral frequencies were tallied across the study period: huddling, proximity, grooming (dam-to-daughter), grooming (daughter-to-dam), aid-for-victim (dam-to-daughter), and aid-for-victim (daughter-to-dam).

As an initial step in determining which behavioral domains should be analyzed separately versus combined, we calculated Pearson correlation coefficients among all candidate outcome variables and key demographic predictors (Table S1). This exploratory assessment quantified covariation among affiliative and supportive behaviors across dam–adult daughter dyads.

Grooming (daughter-to-dam) and grooming (dam-to-daughter) were moderately correlated (*r* = 0.44; Table S1), suggesting that grooming is often reciprocal but sufficiently asymmetric to warrant separate modeling. Grooming also showed moderate positive associations with spatial closeness, such that dyads with higher grooming rates also tended to spend more time huddling or in proximity (e.g., grooming daughter-to-dam with huddling: *r* = 0.43; grooming dam-to-daughter with proximity: *r* = 0.46). Huddling and proximity were highly correlated (*r* = 0.83), indicating substantial redundancy between these spatial measures. Based on this strong overlap, huddling and proximity totals were combined into a composite measure of spatial closeness in subsequent analyses. In contrast, agonistic support (aid-for-victim) showed only weak correlations with grooming and spatial closeness (all *r* values ≤ 0.27), and AFV (dam-to-daughter) and AFV(daughter-to-dam) were also weakly correlated (r = 0.29), suggesting that coalitionary support represents a partially independent component of dam–daughter relationships rather than simply scaling with affiliative contact.

Finally, both dam age and daughter age were negatively associated with affiliative behaviors (e.g., subject age with huddling: *r* = –0.37), consistent with developmental shifts in maternal investment and contact patterns across the adult life span. Together, these correlation patterns supported our analytical approach of treating both directions of grooming and agonistic support as separate behavioral outcomes while combining highly collinear spatial measures, huddling and proximity, into a single closeness variable.

### 2.4 Demographic and Social Predictor Variables

#### 2.4.1 Age Variables

Dam age and daughter age were primary demographic predictors and were examined both as continuous variables (years) and as categorical life-history stages (Table 1). Dam age and daughter age were strongly correlated (r = 0.696), reflecting the biological constraint that dams must be at least three years older than their daughters. Categorical age variables were constructed based on life-history stage boundaries informed by reproductive patterns in this population and by considerations of sample balance across categories.

**Table 1.** Dam and adult daughter age categorization.

|  | **Dam Age Category** | 7.0 – 9.4 years | 10.0 – 13.4 years | 14.0 – 25.5 years |
| --- | --- | --- | --- | --- |
| **Daughter Age Category** |  | **Adult** | **Advanced age** | **Geriatric** |
| 2.9 – 4.4 years | **Subadult** | 23 | 19 | 9 |
| 5.0 – 8.4 years | **Adult** | 8 | 34 | 22 |
| 9.0 – 12.3 years | **Advanced age** | n/a | 4 | 16 |
| 13.0 – 18.2 years | **Geriatric** | n/a | n/a | 6 |

Among daughters, individuals 2.9–4.4 years old at the start of the study (approximately September 1) were classified as ‘subadult’ (N = 51); those 5.0–8.4 years old as ‘adult’ (N = 64); those 9.0–12.3 years old as ‘advanced age’ (N = 20); and those 13.0–18.2 years old as ‘geriatric’ (N = 6). These thresholds reflect known reproductive transitions derived from ENPRC colony production data: females younger than five years show substantially lower reproductive success among 3–4 year old females (10–50% give birth) compared with females aged 5–8 years (70–85%).

Among dams, individuals 7.0–9.4 years old were categorized as ‘adult’ (N = 31), those 10.0–13.4 years old as ‘advanced age’ (N = 57), and those 14.0–25.5 years old as ‘geriatric’ (N = 53). No dams younger than seven years were present in the dataset. The upper boundary for ‘advanced age’ daughters (< 9.0 years) differed from that of dams (< 10.0 years) to maintain adequate representation across cross-classified dam–daughter age combinations and avoid sparsity within analytic cells.

The dam–daughter age gap was calculated as a continuous variable and categorized using two complementary approaches: (1) a three-level classification of smaller, mid-range, and larger age gaps, and (2) a binary classification of mid-range and non-mid-range age gaps. Both were derived from quantile-based breaks of the centered age-difference distribution.

#### 2.4.2 Dominance Rank

Female dominance ranks were calculated using the *Perc* package in R, based on all agonistic interactions with clear winners and losers (Fujii et al., 2015). Adult females were arranged into a linear hierarchy and assigned ordinal ranks.

Because dam and daughter ranks were highly correlated, they were not included simultaneously in the same models. Relative rank difference (dam rank – daughter rank) was used to evaluate hierarchical directionality within dyads.

#### 2.4.3 Family Structure

Family size and structure, for both dam and daughter, were represented by the following variables: total number of adult daughters present in the group and whether they currently had an infant. These predictors were expected to influence how maternal time, attention, and coalitionary investment are distributed.

### 2.5 Conceptual Framework and Analytical Strategy

Dam–daughter relationships involve multiple affiliative behaviors whose direction and relative frequency may vary across life-history stages. We therefore treated grooming, huddling, proximity, and agonistic support as distinct components whose combination defines the behavioral architecture of the relationship rather than as interchangeable indicators of a single latent “bond strength” dimension.

Guided by Hinde’s framework of social relationships as patterned configurations of interaction shaped by both life-history context and the broader social environment (Hinde, 1976), we adopted a two-part analytical strategy designed to address complementary questions. First, we used generalized linear models (GLMs) to identify demographic and social predictors of individual behavioral components of dam–daughter bonds. Second, we used Data Mechanics, an unsupervised multivariate clustering approach (Fushing and Chen, 2014), to identify distinct relationship types defined by distinct combinations of behaviors across life-history stages.

Demographic variables were incorporated according to their hypothesized role in shaping dam–daughter relationships. Absolute ages of dams and daughters were treated as life-history variables defining the developmental context of the dyad and were therefore included in the clustering analysis used to identify relationship types. Variation in dam and daughter age captures differences in maternal life stage and daughter maturity that may influence patterns of maternal investment, reciprocity, and coalitionary support within the dyad.

In contrast, dominance rank, relative rank difference, and family structure variables were expected to influence the expression and distribution of behaviors without altering the underlying kinship role or life-history stage of the relationship. For example, a high-ranking dam with a four-year-old daughter and a low-ranking dam with a four-year-old daughter occupy the same maternal life stage, even though rank may influence the frequency or context of their interactions. These variables were therefore treated as contextual predictors and examined primarily through the GLMs rather than used to define relationship clusters.

### 2.6 Generalized Linear Models of Individual Behaviors

We analyzed grooming and aid-for-victim (AFV) behaviors using generalized linear models for overdispersed count data. Preliminary examination showed that outcome variances substantially exceeded means (variance > mean across all outcomes), so all models were fit using a negative binomial distribution. Directional outcomes were modeled separately (grooming: dam-to-daughter vs. daughter-to-dam; AFV: dam-to-daughter vs. daughter-to-dam).

Because behavioral counts depend on observation effort and dyad availability, all models included an offset term accounting for the number of observation sessions in which both members of a dyad were present. For AFV models, however, the opportunity for support depended on exposure to aggression. Accordingly, dam-to-daughter AFV models included an offset for conflicts in which the daughter was the victim, and daughter-to-dam AFV models included an offset for conflicts in which the dam was the victim.

Because multiple dyads could share the same dam, we evaluated potential non-independence by fitting models that included dam identity as a random intercept. In several cases, the estimated variance of this random effect was effectively zero and models produced singular fits, indicating negligible between-dam variation relative to within-dyad variation. In other cases, the random effect variance was non-zero, but including it did not materially change fixed-effect estimates or their statistical significance. Given these results and to maintain consistency across models, we report results from models without random effects.

Model selection was based on Akaike Information Criterion (AIC), and best-fit models are reported for each behavioral outcome (Burnham and Anderson, 2002).

### 2.6 Data Mechanics: Multivariate Identification of Relationship Types

To examine how affiliative behaviors co-occur across dam–daughter dyads, we applied Data Mechanics, an unsupervised clustering method designed to detect multiscale block structure in behavioral–demographic matrices (Fushing and Chen, 2014). Data Mechanics has been applied across a range of complex data types, including ecological, physiological, and behavioral datasets, to identify structured patterns in multivariate systems (Fushing et al., 2024, 2018, 2015; McCowan et al., 2016). Our goal was to identify recurring types of dam–daughter relationships based on multivariate behavioral profiles while preserving their multidimensional structure.

We selected Data Mechanics because it simultaneously clusters both rows (dyads) and columns (behaviors), allowing the structure of behavioral domains to emerge from the data rather than being imposed a priori (Fushing and Chen, 2014). This was critical, as we were interested not only in variation among dyads, but also in how behavioral domains (e.g., grooming, spatial closeness, agonistic support) organize to define relationship structure.

Rows in the clustering matrix represented dam–daughter dyads, and columns represented behavioral and selected demographic variables. Behavioral variables included huddling, proximity, directional grooming (dam-to-daughter), and directional agonistic support (aid-for-victim from dam to daughter and new-join-with from daughter to dam). These directional measures were selected because they capture developmentally meaningful aspects of the relationship, including maternal investment and daughter participation in coalitionary contexts.

Preliminary analyses including additional variables (e.g., grooming daughter-to-dam, new-join-with dam-to-daughter) produced similar overall structure but reduced clustering clarity, likely due to added noise. We therefore retained a parsimonious set of variables that captured the primary axes of behavioral interaction while maximizing interpretability.

Absolute dam age and daughter age were included as life-history variables defining developmental context. In contrast, dominance rank and family composition were treated as contextual predictors in the GLMs, as they were expected to influence behavioral rates without altering the underlying structure of the relationships.

We transformed all behavioral variables into ordinal categories prior to clustering to enable comparison across measures with markedly different scales and distributions. Raw counts varied widely across behaviors (e.g., mean ± SD: huddle/proximity = 55.97 ± 40.07; grooming = 9.48 ± 9.93; aid-for-victim daughter-to-dam = 0.92 ± 1.67; Table 2), such that using raw values would have caused high-frequency behaviors to dominate the analysis.

**Table 2.** Descriptive statistics for outcome variables.

|  | Mean | SD |
| --- | --- | --- |
| Groom (dam to daughter) | 9.48 | 9.93 |
| Groom (daughter to dam) | 11.31 | 11.88 |
| Huddle & Proximity | 55.97 | 40.07 |
| Aid for victim (dam to daughter) | 3.25 | 3.41 |
| Aid for victim (daughter to dam) | 0.915 | 1.67 |

In addition, equivalent numerical differences do not carry the same behavioral meaning across measures. For example, a change from 0 to 1 instance of agonistic support reflects the presence versus absence of coalitionary behavior, whereas a change from 55 to 56 instances of proximity is unlikely to represent a meaningful shift in the relationship. Converting variables to ordinal categories mitigates these differences in scale and interpretability while preserving relative variation.

We recoded each variable onto a 5-point ordinal scale (from none to high), with thresholds defined separately for each behavior based on its distribution and behavioral interpretation. This approach preserved meaningful distinctions at the low end (e.g., presence versus absence of support) while reducing the influence of small differences at the high end of more frequent behaviors. Dam and daughter age were similarly recoded into ordinal categories to allow inclusion alongside behavioral variables in the clustering framework.

We used the same number of ordinal levels for all variables to ensure comparable contributions to clustering. Variables with more levels can introduce finer-grained structure, whereas those with fewer levels contribute less detail; mixing scales of differing resolution can therefore reduce stability and interpretability. Using a consistent 5-level scale ensured balanced contributions across variables in detecting multivariate structure.

Data Mechanics iteratively reorders rows and columns to minimize an energy criterion, producing a configuration that reveals hierarchical block structure in both dyad and variable space (Fushing and Chen, 2014). In its original formulation, this energy function is derived from an Ising-model framework and reflects the extent to which similar values are grouped into contiguous blocks, with lower energy corresponding to more homogeneous structure across scales.

Because this energy function was meant for use with binary data, it is not directly interpretable for continuous or ordinal data. We therefore calculated within-block variance as a complementary and more transparent measure of block homogeneity. Specifically, within-block variance was computed as the sum of squared deviations of normalized values from their corresponding block means across all row–column cluster intersections (i.e., blocks). Lower within-block variance indicates greater homogeneity within blocks and closer alignment with the low-energy configurations targeted by the algorithm. This metric was subsequently used to compare candidate clustering configurations.

#### Cluster configuration selection

We first evaluated the number of column clusters by comparing configurations with three versus four behavioral clusters across a range of row-cluster resolutions. Although within-block variation did not uniformly favor one option, configurations with four column clusters more consistently produced groupings that aligned with the empirical correlation structure among behaviors. In particular, four clusters preserved distinctions between key behavioral domains (e.g., grooming, spatial proximity, and forms of agonistic support), whereas three-cluster solutions more frequently collapsed these domains into broader groupings, reducing interpretability. Based on this, we fixed the number of column clusters at four.

Then we evaluated candidate row-cluster configurations (6–11) while holding the number of column clusters constant at four. Selection was based on a combination of quantitative and qualitative criteria assessing model fit, structural stability, and biological interpretability.

##### Within-block variation (goodness of fit)

For each configuration, we calculated within-block variation (see above) as a measure of goodness of fit, with lower values indicating more homogeneous block structure.

##### Column clustering validity

We assessed whether column clusters were consistent with the empirical correlation structure among behavioral variables, examining whether highly correlated behaviors (e.g., huddling and proximity, *r* = 0.83) tended to cluster together, and similarly, whether more weakly correlated behaviors (e.g., huddling and grooming, *r* = 0.47) were in separate clusters.

##### Row cluster stability across resolutions

We compared cluster membership across adjacent configurations to evaluate stability. Stable solutions showed consistent membership across resolutions, whereas substantial reorganization indicated under- or over-partitioning.

##### Distinguishing structural vs. within-cluster variation

We evaluated whether increasing the number of clusters revealed qualitatively distinct relationship types or simply subdivided existing clusters along a single dimension. Configurations that introduced additional clusters without changing overall structure were considered over-resolved.

The selected configuration minimized within-block variation while preserving stable cluster membership, maintaining consistency with the correlation structure of the data, and capturing distinct, biologically interpretable relationship types without unnecessary subdivision.

The resulting row clusters represent recurring dam–daughter relationship types, while column clusters identify behavioral domains that co-vary across dyads. Because Data Mechanics is descriptive, it was used to characterize multivariate relationship structure and complement, rather than replace, the hypothesis-driven GLM analyses.

## 3 Results

We present results in two stages, reflecting complementary perspectives on dam–daughter social bonds. First, we report behavior-specific generalized linear models (negative binomial) to identify how demographic variables predict variation in individual behaviors within dam–daughter dyads. Second, we present results from the clustering analysis (Data Mechanics) that integrates these behaviors to identify recurring, multivariate dam–daughter relationship types. Descriptive statistics for all outcome variables are provided in Table 1.

### 3.1 Demographic predictors of individual affiliative & support behaviors

#### 3.1.1 Grooming: dam to daughter

Rates of grooming from dams to daughters were strongly associated with family size, age gap, and dominance relationships (Supplemental Information Table S2). Dams groomed individual daughters less frequently as the number of adult daughters present increased (β = −0.49, p < 0.001), indicating a dilution of maternal grooming effort across multiple adult offspring. Grooming was also higher in dyads with a mid-range age gap (5.6–8.1 years) compared to dyads outside this range (β = 0.33, p = 0.021).

Dominance relationships showed weaker effects. There was a positive trend indicating increased grooming toward daughters who outranked their dams (rank difference; β = 0.04, p = 0.086). In addition, higher-ranking dams groomed daughters more frequently overall (β = −0.76, p = 0.002).

#### 3.1.2 Grooming: daughter to dam

Grooming from daughters to dams was primarily structured by daughters’ life-history stage and family size (Table S3). Daughters groomed their dams less frequently as they aged (β = −0.094, p = 0.004). In addition, daughters with adult offspring of their own groomed their dams at lower rates than daughters without offspring (β = −0.476, p = 0.002).

As in other models, the dams with more adult daughters present showed significantly less grooming per daughter (β = −0.333, p < 0.001). Unlike dam-to-daughter grooming, daughter-to-dam grooming showed no association with dominance rank or rank differences.

#### 3.1.3 Aid-for-victim: dam to daughter

Rates of maternal agonistic support were strongly associated with dam characteristics (Table S4). After controlling for exposure to aggression, the best-fit model showed that older dams were more likely to support their adult daughters (β = 0.075, p < 0.001), and higher-ranking dams provided more support than lower-ranking dams (β = 0.84, p = 0.003). As with grooming, dams with more adult daughters provided less aid per daughter (β = −0.416, p < 0.001).

#### 3.1.4 Aid-for-victim: daughter to dam

Agonistic support directed from daughters to dams was rare (see Table 1) and showed a different demographic pattern (Table S5). After controlling for exposure to aggression, daughters were more likely to support geriatric dams compared to prime-age dams (β = 1.135, p < 0.001). The dam’s total number of adult daughters was again associated with reduced support from any given daughter (β = −0.474, p = 0.001).

#### 3.1.5 Spatial closeness (huddle + proximity)

To evaluate demographic predictors of non-directional affiliation, we modeled a combined measure of huddling and proximity, which were highly correlated (r = 0.83). The best-fit model showed that spatial closeness declined as daughters aged (β = −0.103, p < 0.001).

Spatial closeness was higher in dyads with a mid-range age gap (β = 0.700, p < 0.001), and dams with more adult daughters showed reduced closeness with individual adult daughters (β = −0.151, p = 0.003). An interaction between dam rank and age gap indicated that among dyads with a mid-range age gap, higher-ranking dams showed lower spatial closeness (mid-range age gap × dam rank: β = −0.837, p = 0.012; Table S6).

Across all behaviors, demographic predictors did not exert uniform effects. Dam’s total number of adult daughters present showed a consistent negative relationship with grooming, spatial closeness, and agonistic support, whereas age- and rank-related variables showed behavior-specific effects. This heterogeneity suggests that demographic factors do not simply strengthen or weaken dam–daughter bonds along a single affiliative axis. Instead, a multivariate analysis may be needed to understand how these behaviors are combined within relationships.

### 3.2 Data Mechanics reveals discrete, multidimensional dam–daughter relationship types

We used Data Mechanics clustering to examine the multivariate structure of dam–daughter bonds to identify distinct relationship types. Together, Figures 1–2 and Table 3 provide complementary views of observed dam–daughter relationship structures, from multivariate organization to behavioral expression profiles.

**Figure 1.**
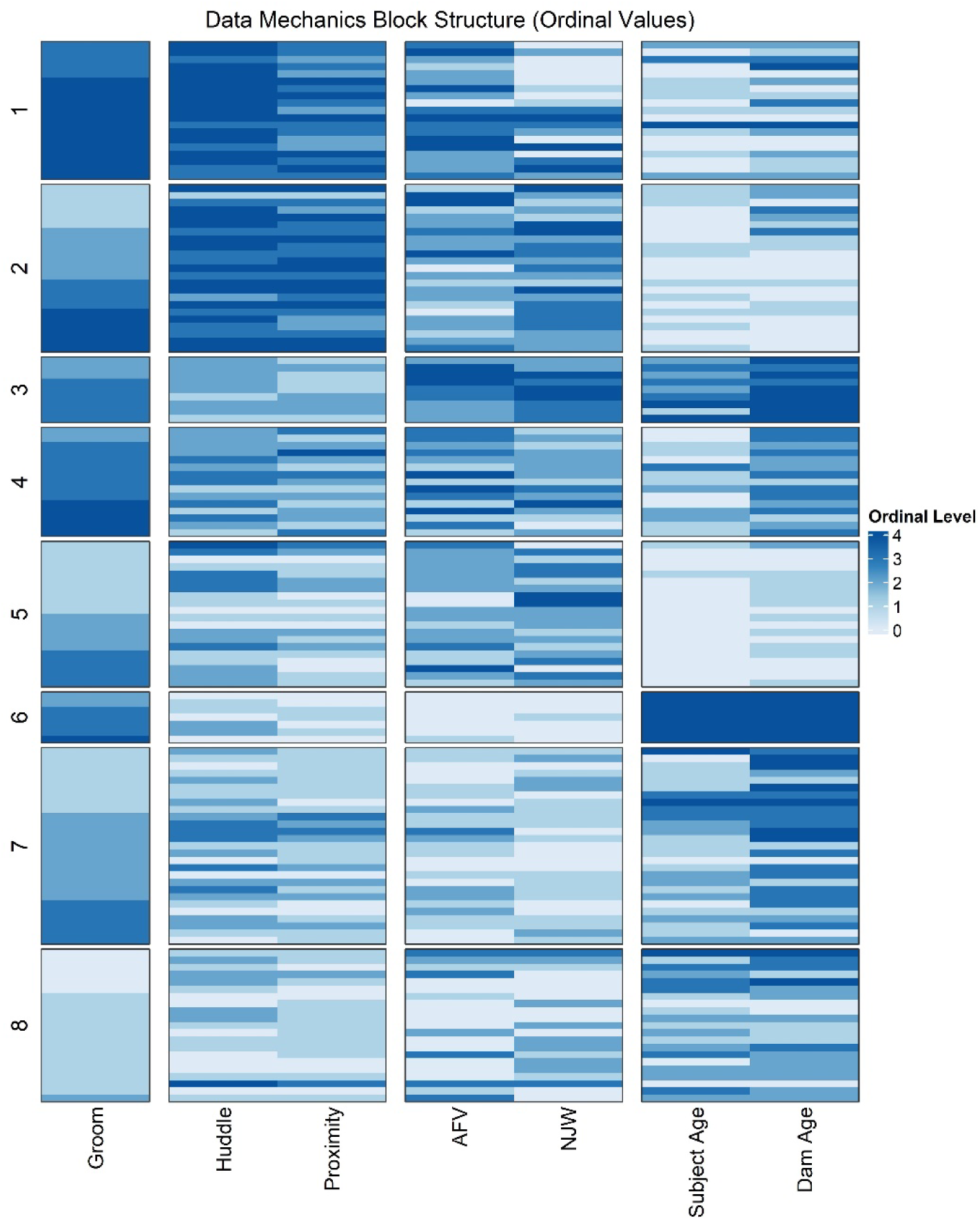
Multidimensional organization of dam–daughter bonds. Heatmap of a dyad-by-variable matrix reordered via iterative clustering of dyads (rows) and variables (columns). Values are transformed to ordinal scales (0–4), with darker shading indicating higher values. The row dendrogram identifies eight clusters corresponding to recurring relationship types, while the column dendrogram organizes variables into behavioral domains, including spatial closeness (huddle, proximity), agonistic support (aid-for-victim, new-join-with), grooming, and age. Clusters differ in both overall behavioral expression and cross-domain configuration, indicating structural variation rather than a single continuum.

**Figure 2.**
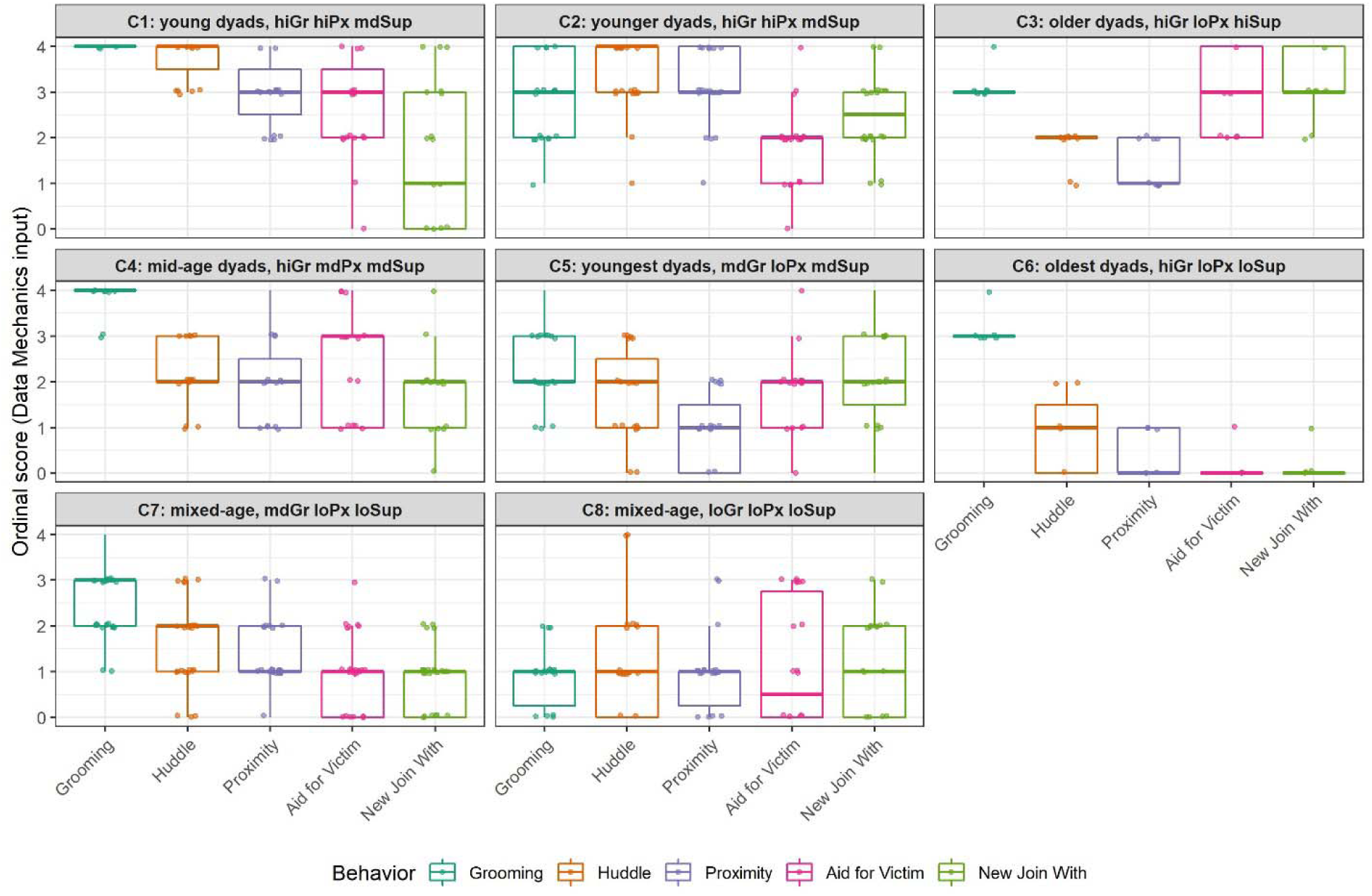
Behavioral profiles of dam–daughter relationship types. Boxplots of ordinal scores (0–4) for grooming, huddling, proximity, aid-for-victim (AFV), and new-join-with (NJW) across the eight dam–daughter relationship types identified by Data Mechanics. Each panel corresponds to one relationship type. Behaviors are ordered along the x-axis to reflect the column clustering structure, with spatial closeness (huddling and proximity) and agonistic support (AFV and NJW) grouped together. Differences among relationship types reflect variation in both the magnitude and configuration of behavioral domains. Panel labels provide abbreviated summaries of cluster characteristics (Gr = grooming, Px = spatial proximity, Sup = agonistic support; hi, md, and lo indicate relative levels).

**Table 3.**
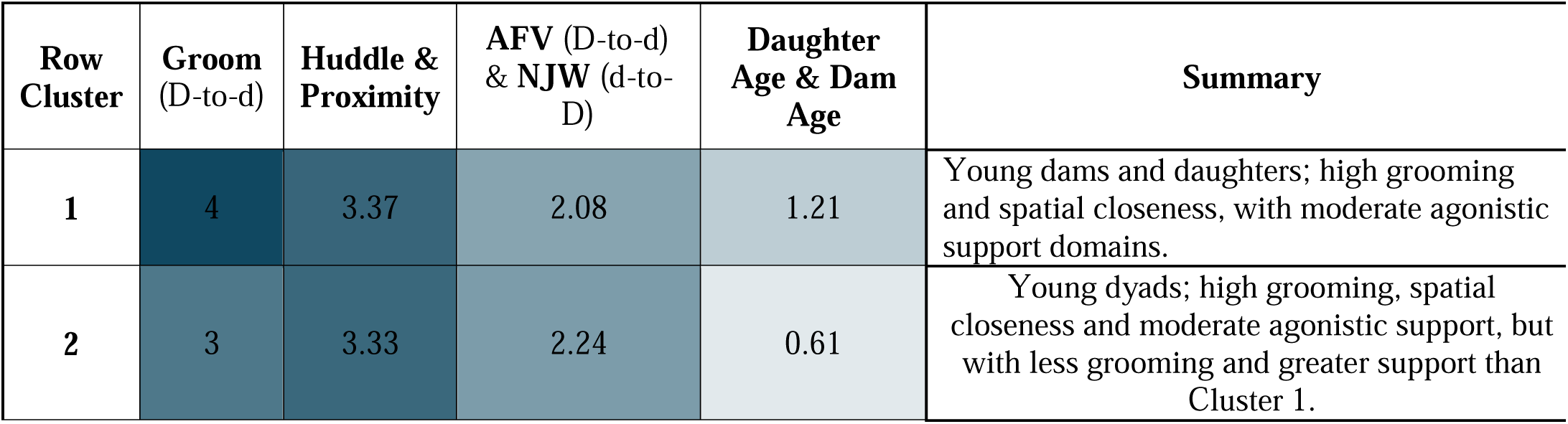
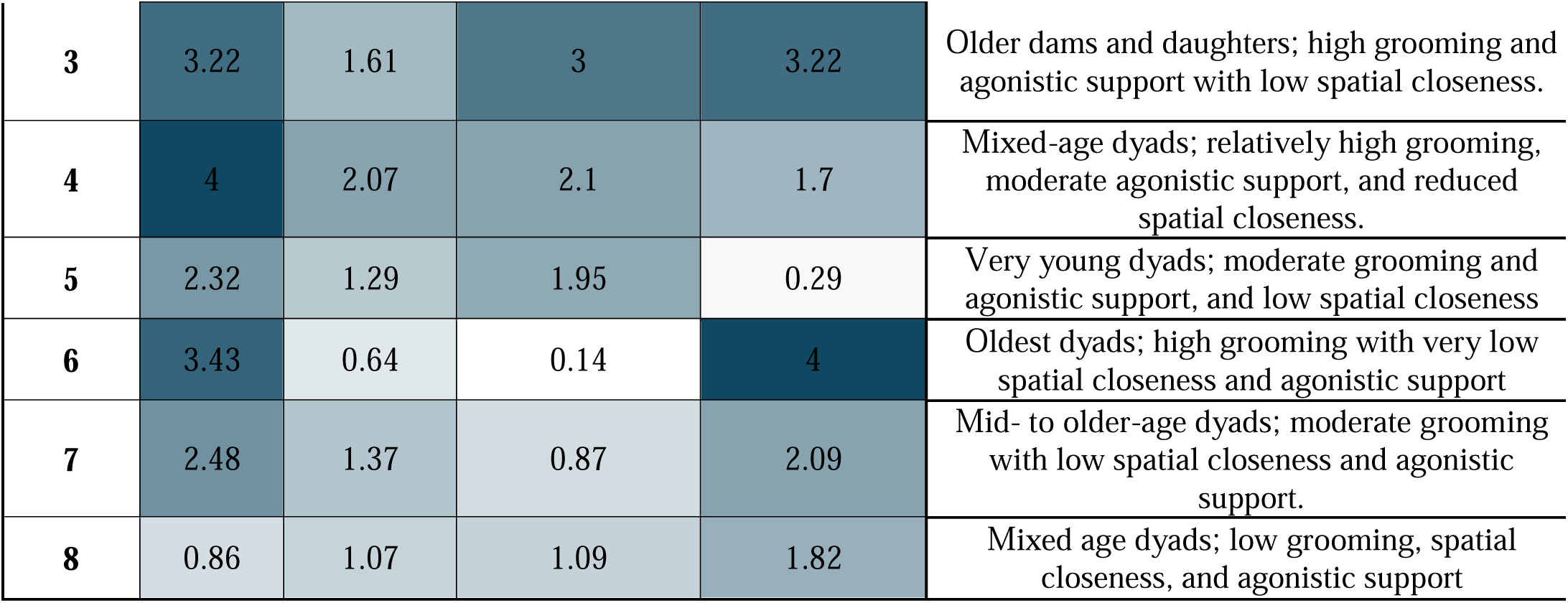
Behavioral and demographic profiles of dam-daughter relationship types. Mean ordinal values (0–4) for behavioral and demographic measures across the eight dam–daughter relationship types identified by Data Mechanics. Values represent cluster-level averages for grooming (dam to daughter), spatial closeness (huddling and proximity), agonistic support (aid-for-victim and new-join-with), and age variables (daughter and dam). Rows correspond to relationship types and columns summarize behavioral and demographic domains. Directional measures are indicated by dam to daughter (D-to-d) or daughter to dam (d-to-D).

| Row Cluster | Groom (D-to-d) | Huddle & Proximity | AFV (D-to-d) & NJW (d-to-D) | Daughter Age & Dam Age | Summary |
| --- | --- | --- | --- | --- | --- |
| 1 | 4 | 3.37 | 2.08 | 1.21 | Young dams and daughters; high grooming and spatial closeness, with moderate agonistic support domains. |
| 2 | 3 | 3.33 | 2.24 | 0.61 | Young dyads; high grooming, spatial closeness and moderate agonistic support, but with less grooming and greater support than Cluster 1. |
| 3 | 3.22 | 1.61 | 3 | 3.22 | Older dams and daughters; high grooming and agonistic support with low spatial closeness. |
| 4 | 4 | 2.07 | 2.1 | 1.7 | Mixed-age dyads; relatively high grooming, moderate agonistic support, and reduced spatial closeness. |
| 5 | 2.32 | 1.29 | 1.95 | 0.29 | Very young dyads; moderate grooming and agonistic support, and low spatial closeness |
| 6 | 3.43 | 0.64 | 0.14 | 4 | Oldest dyads; high grooming with very low spatial closeness and agonistic support |
| 7 | 2.48 | 1.37 | 0.87 | 2.09 | Mid- to older-age dyads; moderate grooming with low spatial closeness and agonistic support. |
| 8 | 0.86 | 1.07 | 1.09 | 1.82 | Mixed age dyads; low grooming, spatial closeness, and agonistic support |

#### 3.2.1 Selection of cluster configuration

Within-block variation decreased from the 6×4 configuration of row and column clusters to 9×4, reaching a local minimum at 9×4, before increasing at 10×4 and 11×4 (Table S7). However, evaluation of cluster structure revealed that solutions differed qualitatively in their interpretability.

The 6×4 configuration produced column clusters that were inconsistent with the correlation structure of the behavioral variables, collapsing grooming with spatial proximity while treating agonistic support behaviors independently. This resulted in row clusters driven disproportionately by individual behaviors and failed to identify low-engagement dyads observed in higher-resolution configurations.

The 7×4, 8×4, and 9×4 configurations showed high stability in row cluster membership, with the 7×4 solution merging two clusters identified in the 8×4 solution (i.e., row clusters 1 and 2), and the 9×4 solution subdividing a single cluster (i.e., row cluster 1) without altering the overall structure.

In contrast, the 10×4 and 11×4 configurations resulted in substantial changes to both column clustering and row cluster membership, producing fragmented clusters driven primarily by variation in agonistic support behaviors AFV and NJW.

Although the 9×4 configuration yielded the lowest within-block variation, the additional cluster reflected subdivision of an existing high-engagement cluster based primarily on differences in agonistic support, rather than the emergence of a new relationship type.

Based on these criteria, the 8×4 configuration was selected as the optimal solution, representing the minimal resolution that captured all major multidimensional relationship types while maintaining stability and interpretability.

#### 3.2.2 Emergent organization of dam–daughter bonds

The Data Mechanics analysis identified eight distinct row clusters, representing recurring dam–daughter relationship types (Figure 1). These clusters captured most of the observed variation, with only three dyads identified as outliers. For additional information about identification of outliers, see Table S7 in Section 4 of the Supplemental Information.

The row dendrogram showed a primary bifurcation separating dyads with broader multi-domain behavioral expression (Clusters 1–5) from those with more limited or uneven expression across domains (Clusters 6–8). Importantly, clusters differed not only in overall levels of behavior, but in the configuration of behaviors across grooming, spatial closeness, and agonistic support (Figure 1, Table 3).

#### 3.2.3 Behavioral Organization Across Domains

Column clustering revealed four major domains: (1) huddle and proximity clustered together, representing spatial closeness; (2) agonistic support measures clustered separately (aid-for-victim from dam to daughter, new-join-with from daughter to dam); (3) dam and daughter age variables; and (4) grooming from dam to daughter as a distinct affiliative axis. The column dendrogram showed that age variables diverged first from the behavioral measures, consistent with the distinction between demographic attributes and social interaction variables. This structure indicates that grooming, spatial closeness, and agonistic support represent partially independent components of dam–daughter bonds.

Notably, certain behavioral combinations were rare. Dyads showing elevated spatial closeness or agonistic support in the absence of grooming occurred in fewer than 10% of cases, indicating that grooming is a foundational component of broadly expressed affiliative relationships.

#### 3.2.4 Relationship Categories and Life-History Stages

Clusters were further characterized based on their position on the ordinal scale across behavioral domains. We identified multi-domain clusters (Clusters 1–5) as dyads exhibiting scores at or above the ordinal scale midpoint (≥2 on a 0–4 scale) in at least two of the three behavioral domains (grooming, spatial closeness, and agonistic support; Table 3). This criterion identifies relationships with moderate or greater expression across multiple behavioral domains, rather than elevated values in a single behavior. Requiring elevated values in at least two domains ensured that classifications reflected multidimensional relationship structure, consistent with our conceptual framework, rather than isolated behavioral tendencies.

Within this set of clusters, multiple relationship profiles emerged. Clusters 1 and 2 were characterized by consistently high expression across grooming and spatial closeness, and moderate agonistic support, and were primarily composed of dyads with relatively young dams and young adult daughters. Cluster 3 included older dams and daughters with high levels of grooming and agonistic support but reduced spatial closeness, indicating a shift toward interaction-based rather than proximity-based expression. Cluster 4 showed relatively high grooming and moderate agonistic support with lower spatial closeness, while Cluster 5 represented dyads with moderate expression across domains that nonetheless met the multi-domain criterion.

Clusters 6–8 were characterized by more limited expression across domains, with dyads exhibiting scores below the ordinal scale midpoint (≤2 on a 0–4 scale) in at least two of the three behavioral domains. Cluster 6 was distinguished by high levels of grooming in the context of low spatial closeness and agonistic support and consisted of the oldest dam–daughter pairs. Clusters 7 and 8 showed progressively reduced expression across domains, with Cluster 8 exhibiting uniformly low levels across all behavioral measures. More detailed descriptions of the characteristics of the row clusters can be found in the Supplemental Information.

### 3.3 Convergence with GLMs

Cluster membership was strongly associated with dam family size, despite the dam’s total number of adult daughters not being included in the clustering. Of the 141 dyads studied, 44 represented dams with only one adult daughter, and 37 of these (84%) were classified into multi-domain bond clusters (1–5) with the largest number (N=13 dyads) in Cluster 1, whereas 29 of 43 (67%) pairs representing dams with three or more adult daughters were classified into limited-domain bond clusters (6–8) with the largest number (N=13 dyads) in Cluster 7. This convergence suggests that family size shapes the range of multivariate relationship types observed, rather than simply lowering individual behavioral frequencies.

Age-related effects observed in the GLMs were similarly clarified by clustering. Older dyads often retained grooming and support while showing reduced spatial closeness (e.g., Cluster 3), explaining why age predicted grooming and proximity in the GLMs but not uniformly agonistic support.

### 3.4 Behavioral Combinations Clarifies Model Structure

The clustering analysis also revealed that not all combinations of affiliative and supportive behaviors are equally represented in the data, further clarifying the structure underlying the heterogeneous GLM results. In particular, dyads showing high levels of spatial closeness or agonistic support in the absence of grooming were rare (< 10% of dyads).

Across the full dataset (N = 141), only nine dyads showed elevated agonistic support (AFV ordinal score ≥ 2) paired with low grooming (ordinal score < 2). Similarly small numbers of dyads showed elevated huddling (N=6) or elevated proximity (N=4) under the same conditions. In total, these patterns represented only 14 unique dyads, 12 of which were classified into Cluster 8, the bond category with the most limited behavioral expression.

Together, these results indicate that grooming is a foundational component of multi-domain affiliative bonds of dams and daughters: high spatial closeness and agonistic support rarely occur in its absence. At the same time, grooming varied more independently across relationship types, particularly among the bond clusters with more limited behavioral expression (clusters 6-8), suggesting that grooming alone is not sufficient to produce the full behavioral profile of a strong dam–daughter bond.

## 4 Discussion

### 4.1 Multidimensional dam–daughter relationships

Dam–daughter relationships represent foundational dyadic bonds within rhesus macaque matrilines, providing the basis for kin familiarity, affiliative bias, and transmission of dominance rank across generations (Berman, 2004; Datta, 1986; Rendall, 2004; Sade, 1967). Consistent with this, our results show that these relationships are structured as multidimensional configurations of behavior rather than along a single continuum of bond strength. We identified eight recurring relationship types defined by distinct combinations of grooming, spatial closeness, and agonistic support. These domains represent complementary components of social interaction, and their variation within dyads gives rise to multiple relationship types. For example, some dyads maintained grooming and agonistic support at or above the midpoint despite relatively low spatial proximity (Cluster 3), whereas others showed consistently high expression across all domains (Clusters 1 and 2). Together, these findings indicate that affiliative behaviors function as partially independent components that combine in multiple ways to produce discrete relationship types.

This multidimensional structure is consistent with Hinde’s (1976) framework, in which relationships are defined by patterns of interaction across behavioral domains rather than by any single measure. Previous multivariate approaches similarly show that social behaviors cluster into underlying dimensions of relationship quality. For example, McFarland and Majolo (2011) found that grooming, proximity, and solicitations for grooming loaded onto a common “value” component, while aggression, tolerance, and agonistic support defined a separate “compatibility” dimension. Similarly, our results show that grooming, spatial closeness, and agonistic support emerge as distinct components of social relationships, but with multiple combinations observed across dyads. Some dyads exhibited configurations in which behaviors that often covary were partially decoupled, such as high grooming and agonistic support in the context of low spatial proximity. These results extend prior work by showing that, rather than varying only along continuous dimensions such as “value” and “compatibility,” relationship components recombine into a discrete set of recurring configurations, with eight types observed rather than a continuous range of theoretically possible combinations.

### 4.2 Demographic structuring of behavioral domains

The generalized linear models help explain how this multidimensional structure arises. Demographic variables showed domain- and direction-specific associations rather than uniform effects across behaviors. Family size emerged as a consistent predictor, with both maternal effort and daughter engagement decreasing as the number of adult daughters increased, suggesting that affiliative interactions are distributed across multiple female offspring.

In contrast, age- and rank-related variables showed more selective effects that varied by behavioral domain. Spatial closeness was highest in dyads with mid-range age gaps (5.6–8.1 years) and declined as the number of adult daughters increased. An interaction between dam rank and age gap indicated that among dyads with mid-range age differences, higher-ranking dams showed reduced spatial closeness. Grooming and agonistic support showed distinct demographic patterns: as daughters aged, they groomed their dams less and showed reduced spatial closeness, whereas older dams provided more agonistic support and received more support when geriatric. Dominance rank influenced maternal grooming and support but had little effect on daughters’ behavior. Together, these results suggest that different components of the relationship reflect distinct demographic processes, such that any single behavioral measure provides only a partial view of dam–daughter bond structure.

This interpretation aligns with evidence that the fitness consequences of sociality depend on dyadic relationship structure rather than aggregate measures. Ellis et al. (2019) showed that survival in female rhesus macaques was predicted by the strength of connections to key partners and by the number of weak ties—both dyadic measures—whereas individual-level measures such as total grooming or proximity were not predictive. These findings highlight that relationships with specific partners carry information not captured by overall interaction rates, reinforcing the importance of dyadic, multidimensional approaches.

### 4.3 Linking demographic processes to relationship types

The relationship types identified by Data Mechanics align closely with these demographic patterns, suggesting that the observed configurations capture meaningful structure rather than artifacts of the clustering procedure. Dyads in smaller families were disproportionately represented among clusters with broader multi-domain expression: 84% of dyads involving dams with only one adult daughter were found in Clusters 1–5. In contrast, 67% of dyads involving dams with three or more adult daughters were concentrated in Clusters 6–8, which were characterized by more limited or uneven expression across behavioral domains. This pattern suggests that family composition is associated with how affiliative behaviors are combined within dyads.

Patterns in the clustering analysis also mirror domain-specific effects identified in the GLMs. Dyads characterized by high levels of agonistic support were concentrated within a subset of relationship types and were associated with older dams, consistent with model results showing increased maternal support with age. For example, Cluster 3, which exhibited the highest levels of agonistic support, was composed primarily of dyads involving older dams and daughters.

A similar correspondence can be seen in grooming. Clusters 1, 2, and 5 consisted of dams and daughters occupying earlier life-history stages, and showed moderate to high levels of grooming, consistent with increased grooming in dyads with mid-range age gaps (5.6–8.1 years) identified in the GLMs. Cluster 6, which consisted of the oldest dams and daughters, also exhibited high grooming despite low expression in other domains, indicating that similar levels of grooming can occur in later life-history stages. Together, these patterns show that moderate to high grooming occurs across multiple relationship types and demographic contexts, even when associated with similar underlying age structure.

More broadly, the GLM and clustering results provide complementary perspectives on relationship structure: the GLMs identify predictors of individual behaviors, while the clustering analysis reveals how those behaviors co-occur within dyads.

### 4.4 Matriline Cohesion and Management Implications

Mother–daughter bonds provide the social scaffolding through which kin become familiar and develop affiliative biases (Berman, 2004; Kapsalis, 2004; Rendall, 2004). Because daughters remain in proximity to their mothers, sisters gain familiarity through shared association with their mother, and daughters also interact with grandmothers and other maternal kin via these connections. Variation in the multidimensionality of dam–daughter bonds therefore may influence the development and stability of other kinship bonds, including sister–sister and grandmother–granddaughter relationships.

However, some variation likely reflects life-history progression. Dyads with multiple adult daughters exhibited more limited or uneven expression across domains (Clusters 7–8), consistent with age-related shifts rather than reduced relational quality. In these cases, the scaffolding for other kin relationships may already have been established earlier, suggesting that limited multidimensionality does not necessarily indicate reduced cohesion. This perspective outlines a potential pathway linking dyadic relationship structure to broader patterns of matrilineal organization.

Dyads in smaller families often exhibited multi-domain expression of affiliation (Clusters 1–5), whereas dyads in larger families more often showed limited or uneven expression (Clusters 6–8). If matrilines are composed of dyads that differ in affiliative configuration, this variation may contribute to differences in matrilineal cohesion and, potentially, group stability. Future research should examine whether variation in dyadic relationship structure generates heterogeneity in matriline cohesion.

From an applied perspective, these findings highlight the importance of moving beyond single behavioral indicators when assessing social relationships in captive groups. Grooming has often been used as an index of relationship strength (Ellis et al., 2019; Silk et al., 2010, 2009), but our results show that it does not fully capture the relationship structure. Dyads with similar levels of grooming can differ substantially in spatial association and agonistic support, indicating that reliance on a single measure may obscure meaningful variation.

Converging evidence supports this interpretation. Vandeleest et al. (2025) showed that health correlates of social connectedness depend on whether relationships span multiple behavioral domains, and Balasubramaniam et al. (2019) found that E. coli transmission risk depends on combined centrality across behavioral networks. Together, these findings indicate that the consequences of social relationships depend on how multiple forms of interaction combine.

A key advantage of this framework is that it allows identification of dyads that deviate from expected multi-domain patterns. Importantly, interpretation may be context-dependent: dyads in larger families more often exhibited limited or uneven expression across behavioral domains, but whether this reflects typical adjustment to family structure or reduced affiliative integration remains unclear. Identifying dyads whose behavioral configurations are uncommon for their demographic context may therefore provide a useful basis for monitoring.

Monitoring a single behavioral domain may be insufficient to identify at-risk relationships, as dyads can show typical levels of one behavior while differing in overall configuration. Considering multiple domains may improve detection of variation relevant to matrilineal cohesion. This dyad-level perspective complements family-level metrics and may support earlier detection of changes in social structure, although links to intra-family social instability requires further study.

## 5 Conclusion

In summary, dam–daughter relationships are best understood as multidimensional configurations of behavior rather than a single continuum of bond strength. This perspective has implications for captive management, as considering multiple behavioral domains may help identify dyads that warrant closer monitoring and support efforts to maintain matrilineal cohesion. Future work should examine how these relationship types relate to longitudinal changes in social structure and their potential utility for understanding social stability in captive rhesus macaque groups.

## Acknowledgments

This project was funded by the following NIH awards: R24 OD030035 to K. Ethun and P51 OD011132 to ENPRC. We thank the animal care, colony management, and veterinary staff at the Emory NPRC for their dedication to the management and welfare of the animals.

## Data Availability Statement

The data that support the findings of this study are available from the corresponding author upon request.

## CRediT authorship contribution statement

**Brianne Beisner**: Conceptualization, Formal Analysis, Methodology, Software, Visualization, Writing – original draft, Writing – review and editing. **Kelly F. Ethun**: Conceptualization, Data Curation, Funding Acquisition, Investigation, Project Administration, Resources, Supervision, Writing – review and editing.

## REFERENCES

Balasubramaniam, K., Beisner, B., Guan, J., Vandeleest, J., Fushing, H., Atwill, E., McCowan, B., 2018. Social network community structure and the contact-mediated sharing of commensal E. coli among captive rhesus macaques (Macaca mulatta). PeerJ 6, e4271. 10.7717/peerj.4271

Balasubramaniam, K., Beisner, B., Hubbard, J., Vandeleest, J., Atwill, E.R., McCowan, B., 2019. Affiliation and Disease Risk: Social Networks Mediate Microbial Transmission among Rhesus Macaques. Animal Behaviour 151, 131–143.

Balasubramaniam, K., Beisner, B., Vandeleest, J., Atwill, E.R., McCowan, B., 2016. Social buffering and contact transmission: network connections have beneficial and detrimental effects on *Shigella* infection risk among captive rhesus macaques. Peer J 4, e2630.

Beisner, B., Jackson, M., Cameron, A., McCowan, B., 2011. Detecting instability in animal social networks: genetic fragmentation is associated with social instability in rhesus macaques. PLoS ONE 6, e16365.

Beisner, B., McCowan, B., Bloomsmith, M.A., Lacefield, L., Ethun, K., 2026. A cross-center comparison of the relationship between matriline fragmentation, grooming cohesion, and agonistic behavior in captive rhesus macaque (Macaca mulatta) social groups. bioRxiv 2026.01.18.700196. 10.64898/2026.01.18.700196

Berman, C., Kapsalis, E., 1999. Development of kin bias among rhesus monkeys: maternal transmission or individual learning? Anim. Behav. 58, 883–894.

Berman, C.M., 2004. Developmental Aspects of Kin Bias in Behavior, in: Chapais, B., Berman, C.M. (Eds.), Kinship and Behavior in Primates. Oxford University Press, pp. 317–346.

Bernstein, I., Gordon, T.P., 1974. The function of aggression in primate societies. Am. Sci. 62, 304–311.

Burnham, K.P., Anderson, D.R., 2002. Model Selection and Multimodel Inference: A Practical Information-Theoretic Approach, Second. ed. Springer-Verlag, New York.

Chapais, B., 1992. Role of alliances in the social inheritance of rank among female primates, in: Harcourt, A.H., de Waal, F.B.M. (Eds.), Coalitions and Alliances in Humans and Other Animals. Oxford University Press, New York, pp. 29–59.

Chapais, B., 1988. Experimental matrilineal inheritance of rank in female Japanese macaques. Anim. Behav. 36, 1025–1037.

Datta, S.B., 1986. The role of alliances in the acquisition of rank, in: Else, J.G., Lee, P.C. (Eds.), Primate Ontogeny, Cognition, and Social Behaviour. Cambridge University Prress, New York, pp. 219–225.

Ehardt, C.L., Bernstein, I., 1986. Matrilineal overthrows in rhesus monkey groups. Int. J. Primatol. 7, 157–181.

Ellis, S., Snyder-Mackler, N., Ruiz-Lambides, A., Platt, M.L., Brent, L.J.N., 2019. Deconstructing sociality: the types of social connections that predict longevity in a group-living primate. Proceedings of the Royal Society B 286, 20191991.

Fujii, K., Jin, J., Shev, A., Beisner, B.A., McCowan, B., Fushing, H., 2015.Perc: Using Percolation and Conductance to find information flow certainty in direct network.

Fushing, H., Chen, C., 2014. Data mechanics and coupling geometry on binary bipartite networks. PLoS ONE 9, e106154.

Fushing, H., Hsueh, C.-H., Heitkamp, C., Matthews, M., Koehl, P., 2015. Unravelling the geometry of data matrices: effects of water stress regimes on winemaking. Journal of the Royal Society Interface 12, 20150753.

Fushing, H., Kao, H.W., Chou, E.P., 2024. Topological Risk-Landscape in Metric-Free Categorical Database. IEEE Access 12, 66296–66318.

Fushing, H., Liu, S.-Y., Hsieh, Y.-C., McCowan, B., 2018. From patterned response dependency to structured covariate dependency: Entropy based categorical-pattern-matching. PLoS ONE 13, e0198253. 10.1371/journal.pone.0198253

Haertel, A.J., Beisner, B., Buehler, M.S., Capuano, S., Carroll, K.E., Church, T., Cohen, J., Crane, M.M., Dutton, J.W., Falkenstein, K.P., Gill, L., Hopper, L.M., Hotchkiss, C.E., Lee, G.H., Malinowski, C., Mendoza, E., Sayers, K., Scorpio, D.G., Stockinger, D., Taylor, J.M., 2023. The impact of housing on birth outcomes in breeding macaque groups across multiple research centers. American Journal of Primatology 85, e23554.

Hannibal, D.L., Bliss-Moreau, E., Vandeleest, J., McCowan, B., Capitanio, J., 2016. Laboratory rhesus macaque social housing and social changes: Implications for research. Am. J. Primatol. 10.1002/ajp.22528

Hinde, R.A., 1976. Interactions, relationships, and social structure. Man 11, 1–17.

Johnston, J., Meeker, T., Ramsey, J.K., Crane, M.M., Cohen, J., Ethun, K., 2020. Utility of Automated Feeding Data to Detect Social Instability in a Captive Breeding Colony of Rhesus Macaques (Macaca mulatta): A Case Study of Intrafamily Aggression. Journal of the American Association for Laboratory Animal Science 59, 46–57.

Kapsalis, E., 2004. Matrilineal kinship and primate behavior, in: Chapais, B., Berman, C.M. (Eds.), Kinship and Behavior in Primates. Oxford University Press, pp. 153–176.

McCowan, B., Beisner, B., Hannibal, D., 2017. Social management of laboratory rhesus macaques housed in large groups using a network approach: A review. Behav. Processes. 10.1016/j.beproc.2017.11.014

McCowan, B., Beisner, B.A., Bliss-Moreau, E., Vandeleest, J.J., Jin, J., Hannibal, D.L., Fushing, H., 2016. Connections matter: social networks and lifespan health in primate translational models. Frontiers in Psychology 7, 00433.

McFarland, R., Majolo, B., 2011. Exploring the Components, Asymmetry and Distribution of Relationship Quality in Wild Barbary Macaques (Macaca sylvanus). PLoS ONE 6, e28826. doi:10.1371/journal.pone.0028826

Oates-O’Brien, R.S., Farver, T.B., Anderson-Vicino, K.C., McCowan, B., Lerche, N.W., 2010. Predictors of matrilineal overthrows in large captive breeding groups of rhesus macaques (*Macaca mulatta*). Journal of the American Association for Laboratory Animal Science 49, 196–201.

Rendall, D., 2004. “Recognizing” Kin: Mechanisms, Media, Minds, Modules, and Muddles, in: Kinship and Behavior in Primates. Oxford University Press, Oxford, pp. 295–316.

Rox, A., Waasdorp, S., Sterck, E.H.M., Langermans, J.A.M., Louwerse, A.L., 2022. Multigenerational Social Housing and Group-Rearing Enhance Female Reproductive Success in Captive Rhesus Macaques (Macaca mulatta). Biology 11, 970. 10.3390/biology11070970

Sade, D.S., 1972. A longitudinal study of social behavior of rhesus monkeys, in: Tuttle, R. (Ed.), The Functional and Evolutionary Biology of Primates. Aldine Atherton, New York, pp. 378–398.

Sade, D.S., 1967. Determinants of dominance in a group of free-ranging rhesus monkeys, in: Altmann, S.A. (Ed.), Social Communication Among Primates. University of Chicago Press, Chicago.

Silk, J.B., Alberts, S.C., Altmann, J., 2003. Social Bonds of Female Baboons Enhance Infant Survival. Science 302, 1231–1234. 10.1126/science.1088580

Silk, J.B., Beehner, J.C., Bergman, T.J., Crockford, C., Engh, A.L., Moscovice, L.R., Wittig, R.M., Seyfarth, R.M., Cheney, D.L., 2010. Strong and consistent social bonds enhance the longevity of female baboons. Curr Biol 20, 1359–61. 10.1016/j.cub.2010.05.067

Silk, J.B., Beehner, J.C., Bergman, T.J., Crockford, C., Engh, A.L., Moscovice, L.R., Wittig, R.M., Seyfarth, R.M., Cheney, D.L., 2009. The benefits of social capital: close social bonds among female baboons enhance offspring survival. Proceedings. Biological sciences 276, 3099–104. 10.1098/rspb.2009.0681

Vandeleest, J.J., Wooddell, L.J., Beisner, B., Nathman, A., McCowan, B., 2025. Differential effects of multiplex and uniplex affiliative relationships on biomarkers of inflammation. PeerJ 24, 319113.

