## Supplemental Information for "Structure and Variation in Dam–Daughter Bonds in Captive Rhesus Macaques"

This Supplemental Information provides additional methodological and descriptive detail supporting the Data Mechanics clustering analysis of dam–daughter affiliative relationships as well as the output from the best-fit models from the GLMs fitted to each behavioral outcome variable.

**Section 1. Pearson correlation of behavioral outcome variables and age variables**

**Table S1. Correlation Matrix**

|  | **Daughter Age** | **Dam Age** | **Groom Daughter-Dam** | **Groom Dam-Daughter** | **Huddle Total** | **Prox Total** | **AFV Daughter-to-Dam** | **AFV Dam-to-Daughter** |
| --- | --- | --- | --- | --- | --- | --- | --- | --- |
| **Daughter Age** | 1.00 | 0.70 | -0.37 | -0.12 | -0.37 | -0.35 | -0.06 | -0.17 |
| **Dam Age** | 0.70 | 1.00 | -0.30 | -0.16 | -0.33 | -0.34 | -0.01 | -0.11 |
| **Groom Daughter-to-Dam** | -0.37 | -0.30 | 1.00 | 0.44 | 0.43 | 0.39 | 0.14 | 0.24 |
| **Groom Dam-to-Daughter** | -0.12 | -0.16 | 0.44 | 1.00 | 0.47 | 0.46 | 0.23 | 0.28 |
| **Huddle Total** | -0.37 | -0.33 | 0.43 | 0.47 | 1.00 | 0.82 | 0.08 | 0.25 |
| **Prox Total** | -0.35 | -0.34 | 0.39 | 0.46 | 0.82 | 1.00 | 0.07 | 0.27 |
| **AFV Daughter-to-Dam** | -0.06 | -0.01 | 0.14 | 0.23 | 0.08 | 0.07 | 1.00 | 0.29 |
| **AFV Dam-to-Daughter** | -0.17 | -0.11 | 0.24 | 0.28 | 0.25 | 0.27 | 0.29 | 1.00 |

**Section 2. GLM best-fit models for each behavioral outcome variable**

Here we provide tables of the model outputs from the best-fit GLMs of grooming, huddle-proximity, and aid-for-victim behaviors.

**Table S2.** Best-fit model of grooming from dam to daughter (dAIC = 1.6 compared to next best-fit model, which omitted the Rank Difference variable)

|  | Coefficient | P-value |
| --- | --- | --- |
| Age Gap (mid-range vs non-mid-range) | 0.327 | 0.021 |
| Rank Difference | 0.040 | 0.086 |
| Rank of Dam (% of others she outranks) | -0.757 | 0.002 |
| Dam’s total adult daughters | -0.498 | <0.001 |

**Table S3.** Best-fit model of grooming from daughter to dam (dAIC = 2.3 compared to next best-fit model, which included a non-significant term for dam age)

|  | Coefficient | P-value |
| --- | --- | --- |
| Daughter’s total adult female offspring | -0.476 | 0.002 |
| Daughter age (years) | -0.094 | 0.004 |
| Dam’s total adult daughters | -0.333 | <0.001 |

**Table S4**. Best-fit model of aid-for-victim from dam to daughter

|  | Coefficient | P-value |
| --- | --- | --- |
| Dam age (years) | 0.075 | < 0.001 |
| Rank of Dam (% of others she outranks) | 0.84 | 0.003 |
| Dam’s total adult daughters | -0.416 | < 0.001 |

**Table S5**. Best-fit model of aid-for-victim from daughter to dam

|  | Coefficient | P-value |
| --- | --- | --- |
| Dam age (advanced age vs adult) | 0.277 | 0.36 |
| Dam age (geriatric vs adult) | 1.135 | < 0.001 |
| Dam’s total adult daughters | -0.474 | 0.001 |

**Table S6**. Best-fit model of spatial closeness (huddle + proximity) between dams and daughters

|  | Coefficient | P-value |
| --- | --- | --- |
| Daughter Age (years) | -0.103 | < 0.001 |
| Age Gap (mid-range vs non-mid-range) | 0.700 | < 0.001 |
| Rank of Dam (% of others she outranks) | -0.276 | 0.153 |
| Daughter has current infant | 0.315 | < 0.001 |
| Dam’s total adult daughters | -0.151 | 0.003 |
| Age Gap (mid-range) × Rank of Dam | -0.837 | 0.012 |

**Section 3:**

**Selection of Cluster Configuration**

**Table S7.** Summary of metrics used to evaluate different row and cluster configurations in the clustering analysis of dam-daughter relationship types. Within-block variation measures the level of homogeneity within each block, to quantify goodness-of-fit for each configuration. Column clustering structure notes which behaviors clustered together. Row cluster stability reflects consistency across neighboring configurations (e.g., 7 × 4 and 8 × 4). Behavioral consistency indicates whether behaviors that cluster together (or separately) are in agreement with observed correlations, similar to Column Clustering Structure.

| **Configuration (row** × **column cluster)** | **Within-Block Variation** | **Column Clustering Structure** | **Behavioral Consistency** | **Row Cluster Stability** | **Structural Interpretation** |
| --- | --- | --- | --- | --- | --- |
| 6 × 4 | 584.9 | Groom + spatial merged; AFV, NJW separate | Inconsistent with correlations (groom ≠ spatial) | Unstable (differs from 7 × 4 cluster membership) | Distorted (feature imbalance) |
| 7 × 4 | 491.1 | Groom separate; spatial clustered | Consistent | Differs from 6 × 4, but matches both 8×4 & 9 × 4 membership, except cluster merges) | Under-resolved (merges clusters) |
| 8 × 4 | 467.5 | Groom separate; spatial clustered | Consistent | Stable (matches 7×4 membership, except splits cluster 3) | Optimal |
| 9 × 4 | 450.6 | Same as 8×4 | Consistent | Stable (matches 8×4 membership, except splits cluster 1) | Over-resolved (subdivision only) |
| 10 × 4 | 473.2 | Groom + spatial merged | Inconsistent | Unstable (differs from 8×4 & 9 × 4 membership) | Structural shift in clustering |
| 11 × 4 | 458.3 | Groom + spatial merged | Inconsistent | Intermediate (similar to 10 × 4 membership and 6×4, but differs from 8×4 & 9 × 4 membership) | Degenerate refinement |

**Figure S1.** Stability of row cluster membership across clustering resolutions.
Alluvial plot showing how individual dyads are reassigned across successive Data Mechanics configurations (6×4 through 11×4). Each flow represents a dyad, and colors indicate cluster membership in the 8×4 configuration. Coherent splits (e.g., from 8×4 to 9×4) reflect structured differentiation of relationship types, whereas diffuse fragmentation at higher resolutions indicates over-partitioning.

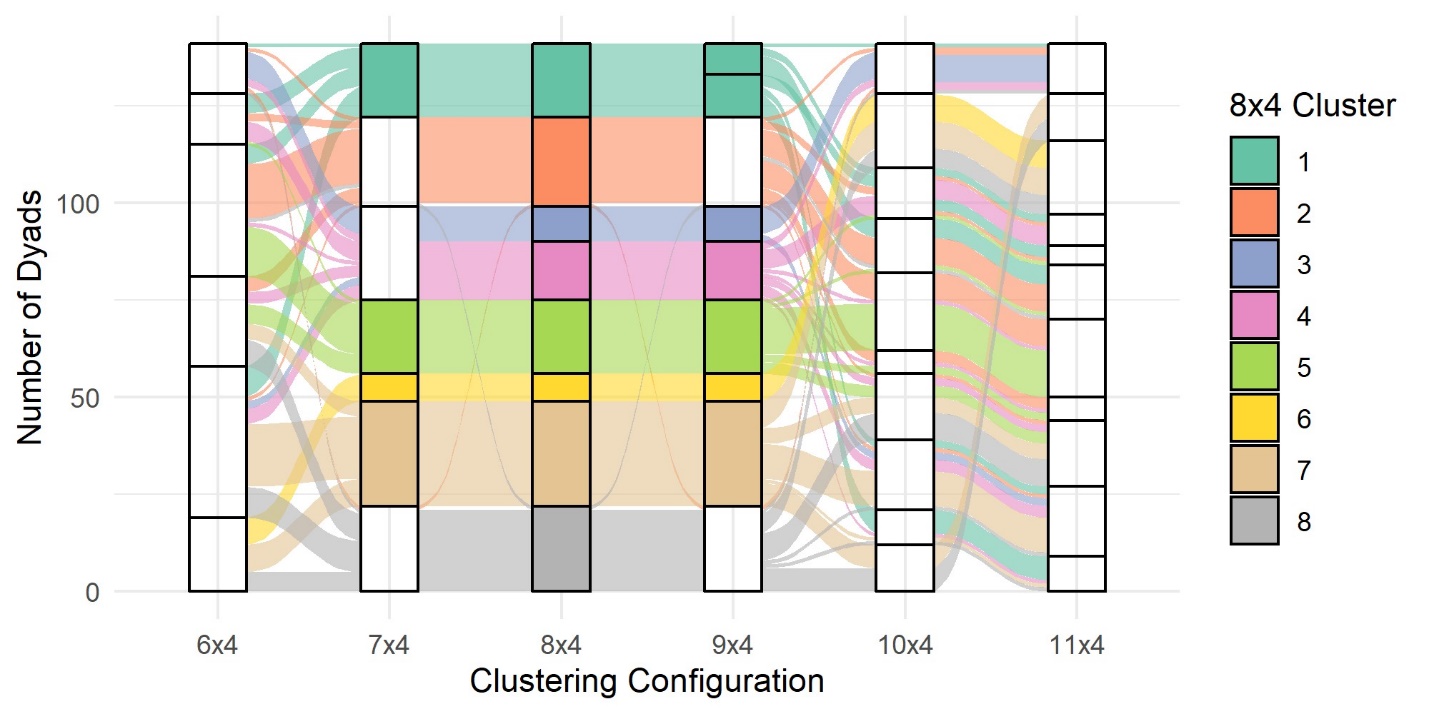

**Additional Details on Relationship-Type Clustering**

Here, we provide (i) the identification and interpretation of potential outlier dyads, (ii) variation in the distribution of relationship types across study groups, and (iii) behavioral and demographic interpretation of each row cluster.

**Behavioral Interpretation of Row Clusters**

Row clustering revealed eight distinct dam–daughter relationship types, spanning a continuum from multidimensional bonds with high affiliative exchange across multiple behavioral domains to more bonds with limited expression of affiliation, characterized by uniformly low interaction rates. Cluster membership was also strongly associated with demographic variables not used in clustering, confirming that the emergent typology reflects meaningful variation in female life history.

**Row Clusters 1–5: Multi-domain affiliative profiles**

**Clusters 1–2: Young dyads with broad affiliative expression**
Dyads in Clusters 1 and 2 consist of young daughters and relatively young dams (Cluster 1: mean daughter age = 5.9 years, mean dam age = 11.9 years; Cluster 2: mean daughter age = 4.4 years, mean dam age = 10.5 years). These pairs exhibit high levels of grooming and spatial closeness (huddling and proximity), along with moderate agonistic support. Together, these clusters represent dyads with consistently high expression across multiple behavioral domains. Cluster 2 differs from Cluster 1 primarily in showing slightly lower grooming intensity.

**Cluster 3: Older dyads with grooming- and support-focused profiles**
Cluster 3 dyads involve substantially older dams and adult daughters (mean daughter age = 9.1 years; mean dam age = 19.0 years). These pairs show high grooming and agonistic support but low spatial closeness, indicating that affiliative relationships are maintained primarily through active social interactions rather than sustained proximity.

**Cluster 4: Mixed-age dyads with differentiated affiliative expression**
Dyads in Cluster 4 show relatively high grooming and moderate agonistic support but reduced huddling and proximity. These relationships fall within the multi-domain affiliative set but are characterized by less spatial cohesion than Clusters 1–3. Cluster 4 dyads include relatively young daughters (mean = 5.8 years) paired with older dams (mean = 14.3 years).

**Cluster 5: Young dyads with moderate multi-domain expression**
Cluster 5 contains the youngest daughters overall (mean = 3.9 years) and young dams (mean = 9.7 years). These dyads show moderate levels of grooming, agonistic support, and spatial closeness. Although lower in magnitude than Clusters 1–2, their behavioral profiles meet the multi-domain criterion and remain distinct from clusters with more limited expression across domains.

**Row Clusters 6–8: Limited or uneven affiliative profiles**

**Cluster 6: Older dyads with grooming-dominant profiles**
Dyads in Cluster 6 include the oldest dams and daughters and show very low spatial closeness and agonistic support. Notably, grooming remains moderate to high, indicating that grooming can persist even when other forms of affiliative exchange are minimal. This cluster represents a pattern in which affiliation is concentrated within a single behavioral domain.

**Cluster 7: Mid- to older-age dyads with low multi-domain expression**
Cluster 7 dyads show moderate grooming, low spatial closeness, and very low agonistic support. These pairs exhibit limited expression across domains, with somewhat greater spatial proximity than Cluster 6 but reduced grooming intensity.

**Cluster 8: Dyads with uniformly low expression across domains**
Cluster 8 represents dyads with low grooming, low spatial closeness, and low agonistic support. Their behavioral profiles reflect minimal affiliative exchange across domains.

**Demographic validation of cluster structure**
Importantly, cluster membership was strongly associated with a demographic variable not included in the clustering procedure: the dam’s total number of adult daughters present in the group. This strongly suggests that the emergent relationship categories reflect meaningful variation in females’ life-history and social context, rather than being an artifact of behavioral measurement alone.

**Section 4. Interpretation of Potential Outlier Dyads**

To evaluate the stability and interpretability of the clustering solution, dyads identified as potential multivariate outliers were examined individually. Outliers were initially flagged based on unusually large Euclidean distances from other dyads within their assigned row cluster. Each flagged dyad was then evaluated qualitatively to determine whether it represented: (1) a true outlier that did not clearly fit any relationship type, (2) a mild outlier that generally matched its assigned cluster but differed in one or two behavioral dimensions, or (3) a misassigned dyad whose behavioral profile more closely matched a different cluster.

Of the eight dyads initially flagged, two were ultimately determined not to be outliers, two appeared to fit a different row cluster more closely, and one was classified as a mild outlier. The remaining three dyads were identified as true outliers, reflecting uncommon combinations of affiliative behaviors not well captured by the eight dominant relationship categories.

One true outlier, D15, was assigned to Cluster 1 and exhibited the high grooming, spatial closeness, and agonistic support characteristic of that relationship type. However, both the dam and daughter were substantially older than is typical for Cluster 1 dyads, making this relationship demographically unusual relative to the rest of the cluster.

A second true outlier, C16, was assigned to Cluster 7 and generally matched that cluster’s pattern of moderate grooming and low new-join-with (NJW). However, this dyad also showed unusually high agonistic support and spatial closeness, producing a behavioral profile that partially overlapped with relationship types characterized by greater multidimensional affiliative engagement. Notably, C16 exhibited higher spatial closeness than grooming, a pattern that is relatively uncommon among mother–daughter dyads and occurs in only ~20% of dyads in this dataset.

Finally, C12 diverged from Cluster 8 by exhibiting unusually high agonistic support despite low grooming and low spatial proximity. This configuration—agonistic support exceeding both grooming and spatial closeness—is also uncommon, occurring in only ~25% of dyads overall, and is disproportionately represented within Cluster 8.

Together, these findings suggest that the clustering solution captures the dominant forms of dam–daughter relationship structure in the dataset, while a small number of dyads exhibit uncommon combinations of affiliative behaviors that fall outside the primary typology.

**Supplementary Table S7** summarizes all potential outlier dyads, their assigned cluster, best-matched cluster (if applicable), and final classification.

**Table S7**. Potential outlier dyads and their official classification (Not an outlier, misassigned, mild outlier, and true outlier), their assigned cluster and the best-matched cluster (if any). Three dyads (D15, C16, C12) were found to be true outliers that did not fit into any of the row clusters identified by Data Mechanics.

| **ID** | **Original Cluster** | **Best Matching Cluster** | **Classification** |
| --- | --- | --- | --- |
| **D15** | 1 | None; maybe 3 or 6 | **True outlier**. Behaviorally consistent with Cluster 1 (high affiliation, very high grooming), but demographically atypical due to unusually old daughter and dam |
| **E20** | 2 | 2 | Not an outlier. Fits Cluster 2 behavioral profile (high affiliation; moderate–high AFV/NJW) but has a slightly older dam than typical for this cluster. |
| **C19** | 4 | 1 | Misassigned. Mostly behaviorally consistent with Cluster 4 (high grooming; moderate AFV; but much higher NJW than Cluster 4) but differs demographically in having a younger-than-typical daughter. |
| **C20** | 5 | 5 | Mild outlier. Matches the Cluster 5 behavioral pattern (lower affiliation; higher grooming) but is unusual due to its very high AFV. |
| **A20** | 7 | 7 | Not an outlier. Fits Cluster 7 behavioral profile (low AFV/NJW; moderate grooming) but differs demographically due to very young dam. |
| **C16** | 7 | None; maybe 3 | **True outlier**. Fits the overall Cluster 7 pattern (low NJW; moderate grooming) but is atypical because both individuals are older than typical for this cluster, with unusually high AFV, huddle, and proximity values. |
| **E21** | 8 | 1 | Misassigned. Demographically more similar to very young dyads like Cluster 1, and behaviorally aligns better with Cluster 1 due to higher values than the Cluster 8 norm on huddle, proximity, AFV and NJW. |
| **C12** | 8 | None; maybe 3 | **True outlier**. Demographically matches older dyads, such as Cluster 3 or 4, and behaviorally diverges from Cluster 8 due to unusually high AFV/NJW for this low-affiliation cluster. |

**Section 5. Distribution of Relationship Types Across Study Groups**

The relative frequency of dam–daughter relationship categories varied substantially across study groups (Supplementary Table S2). Three broad patterns are notable.

First, **Groups A and B** exhibited the highest frequency of multidimensional bond types, with **65%** and **61%** of dyads falling in Clusters 1–4, respectively. In both groups, the most common category was Cluster 2 (**Group A**: 24.2%; **Group B**: 35.3%), representing the most multidimensional dam–daughter bonds observed in the dataset.

Second, **Groups C and D** showed the highest frequency of bonds of limited dimensionality, with **52%** and **56%** of dyads falling in Clusters 6–8, respectively. The most numerous category in Group C was Cluster 8 (29.6%), representing the bonds with the most limited affiliative expression in the dataset, whereas the most numerous category in Group D was Cluster 7 (23.5%), representing the relationship type with next most limited affiliative expression.

Finally, **Group E** showed a somewhat distinct pattern, with the most common dyad category being Cluster 5 and Cluster 7, suggesting a broader distribution across intermediate relationship types.

**Supplementary Table S8** provides the full distribution of dyads across clusters for each study group.

**Table S8. Percentage of dam–daughter dyads per row cluster across study groups**

| **Row Cluster** | **A** | **B** | **C** | **D** | **E** |
| --- | --- | --- | --- | --- | --- |
| 1 | 9.1% | 11.8% | 18.5% | 20.6% | 6.7% |
| 2 | 24.2% | 35.3% | 7.4% | 11.8% | 10.0% |
| 3 | 15.2% | 5.9% | 0.0% | 2.9% | 6.7% |
| 4 | 12.1% | 11.8% | 11.1% | 5.9% | 13.3% |
| 5 | 9.1% | 23.5% | 11.1% | 2.9% | 26.7% |
| 6 | 3.0% | 0.0% | 0.0% | 17.6% | 0.0% |
| 7 | 9.1% | 11.8% | 22.2% | 23.5% | 26.7% |
| 8 | 18.2% | 0.0% | 29.6% | 14.7% | 10.0% |
